# Multiple genetic origins of non-native, self-sustaining rainbow trout *Oncorhynchus mykiss* in streams in Baden-Württemberg, Germany

**DOI:** 10.64898/2025.12.11.693811

**Authors:** J. Peter Koene, Arne Jacobs, Patrick Bartolin, Jan Baer, David Frei, Pascal Vonlanthen, Alexander Brinker

## Abstract

Introduced in the late 19^th^ Century, *Oncorhynchus mykiss* (rainbow trout) have been stocked historically in streams throughout Baden-Württemberg, Germany, and some populations have become self-sustaining with unclear impact on native salmonid populations. We sampled 223 rainbow trout from 14 streams and 3 hatcheries, from which the streams are known to have been stocked. We conducted genomic analyses to uncover evidence confirming self-sustaining populations, to deduce potential sources of these populations, to compare the genetic diversity of hatchery *vs* stream populations, and to discover genetic differences between stream and hatchery populations. We found genetic population structuring amongst the stream populations, consistent with natural reproduction over several generations, and we inferred multiple genetic origins, potentially including source populations beyond the three hatcheries considered. We found no significant difference in genetic diversity between stream and hatchery populations, but there were specific positions in the genome associated with naturalisation within or adjacent to immunity, growth and development genes. Whether such genes are under selection in wild stream environments needs still to be determined to inform fisheries and conservation management.

## Introduction

Since the late 19^th^ Century, *Oncorhynchus mykiss* (Walbaum 1792) (rainbow trout), has been one of the most commonly introduced fish species worldwide (Halverson, 2010). Both its popularity as a game fish and its favourable attributes for aquacultural food production – easy husbandry, tolerance of higher temperatures relative to other native salmonid species, easy manipulation of life history via selective breeding, to name a few – make it economically valuable (Crawford & Muir, 2008; Stanković et al. 2015). Ultimately originating with a small number of source lineages from California in the 1870s, hatchery rainbow trout have been introduced to almost 100 countries, more than half of which now host established self-sustaining populations in wild ecosystems (Stanković et al. 2015). Hatchery fish may be introduced to wild systems through deliberate stocking or as aquaculture escapees (Fausch, 2007). When they naturalise (*i.e.* establish self-sustaining populations) in the wild, they may become potentially invasive, even to the extent of inclusion as a fish example in the IUCN list of the 100 of the world’s worst invasive alien species (Lowe, et al., 2000). Their detrimental effects on native fish fauna in some regions are well documented: among others, predation upon native salmonids, competition for space and resources, redd superimposition, and the introduction and transfer of novel pathogens (reviewed by Stanković et al. 2015). Interestingly, both rainbow trout and brown trout (*Salmo trutta*, L.) may be invasive when introduced to the native areas of other species, with brown trout displaying higher invasiveness and greater negative potential (Gatz et al., 1987; Young et al., 2010). In places such as New Zealand, where both species are non-native, brown trout appear to be more successful, even though contradictory spawning competition there would seem to favour the rainbow trout (Hayes, 1987). Generally, research is lacking on inverse competition in both species and the occurrence of rainbow trout in areas where brown trout are native (Fausch, 2007; McGlade et al., 2022; Stanković et al. 2015).

However, serious declines in native European salmonid populations since the late 20^th^ Century have led to questioning the prudence of rainbow trout stocking (Arlinghaus et al., 2002; Burkhardt-Holm et al., 2002; Stanković et al. 2015), especially as stressors to native salmonids are exacerbated by effects of climate change (Ros et al., 2022; Rubin et al., 2022). In the federal state of Baden-Württemberg in southern Germany, changes in management strategy have seen stocking of rainbow trout decline markedly and even be banned in some catchments to avoid competition with native brown trout (Baer & Brinker, 2010). These bans were imposed in the 1990s as a precautionary measure, based mostly on findings from countries where neither brown nor rainbow trout is native (Hayes, 1987). Yet, populations of rainbow trout persist in a small proportion of formerly stocked streams throughout the state (Stanković et al. 2015). It is known that stocked trout originate from at least three commercial fish farms: Fischzucht Hofer Forellen (with a long history of domestication and advanced technological integration), Fischzucht Störk (medium history of domestication and medium degree of technological integration), and Fischzucht Lohmühle (short history of domestication and low degree of technological integration). However, due to a paucity of records, it is not known from which of these farms the present naturalised populations stem. It is, therefore, also unknown whether descendants from all three hatchery lineages were able to establish themselves successfully in wild streams, or whether specific lineages were more likely than other lineages to naturalise and hence represent a greater potential danger to local fish fauna.

Naturalisation success of rainbow trout beyond its native range is thought to be more dependent upon genetic characteristics than environmental conditions (Koutsikos et al., 2019). It is not known, however, whether parallel naturalisation events resulted from specific highly invasive genetic lineages, or whether they proceed from multiple genomic origins which led to similar ends (Elmer & Meyer, 2011). It has been suggested that multiple phenotypic traits known to have a strong genetic basis may play an important role in naturalisation success (Mueller et al., 2024): spawning time and egg size (Weber et al., 2023); ancestral anadromy *vs* riverine residency, which is underpinned by inversion in a large region of one chromosome, Omy5 (Pearse et al., 2014; 2019); and behavioural traits such as aggressiveness and activity level in the presence of native species (McGlade et al., 2022). Previous work on naturalised populations across European streams using mitochondrial DNA found high levels of genetic diversity compared to hatchery populations, which may be crucial for naturalisation success (Stanković et al. 2016). However, to our knowledge, the greater resolution of interpopulation relationships between self-sustaining rainbow trout and their aquacultural sources in Europe offered by studies that span the genome have not yet been conducted.

We analysed population structure across hatchery and stream populations in southern Germany through restriction-site-associated sequencing (RADseq) of genomic DNA to determine whether genomic data is consistent with natural reproduction across the study streams, to deduce the sources of naturalised populations, to compare the genetic diversity of hatchery *vs* stream populations, and to discover genetic differences, including regions of genomic differentiation, between stream and hatchery populations.

## Methods

### Sampling, sequencing and data processing

The database of the Fisheries Research Station of Beden-Württemberg, which contains data on fish stock surveys in the waters of the German federal state of Baden-Württemberg from the past 40 years (< 80,000 datasets), was searched for potential self-reproducing rainbow trout stocks. In addition, local experts were asked whether they knew of any further stocks of naturally reproducing rainbow trout stock. Subsequently, rainbow trout specimens (*n* = 219) from proven self-sustaining populations in 14 wild streams across Baden-Württemberg were captured by a combination of electrofishing (single anode, 600 V DC electrofishing setup, 0.65 kW, Bretschneider Spezialelektronik) and angling (fly fishing, dry fly or nymph) (Supplementary material, Fig. S1). All fish were caught by licenced personnel under permission of the local fisheries administration (Regierungspräsidium Karlsruhe, Freiburg, Tübingen) according to the German Animal Protection Law (§ 4). Captured fish were killed according to the ordinance on slaughter and killing of animals (Tierschutzschlachtverordnung § 13). Additionally, specimens were sampled from three aquaculture facilities that had previously been used to stock the sampled streams. Those rainbow trout have undergone domestication for up to a century, almost exclusively within their own populations. Fin clips were immediately taken from all specimens after euthanasia, stored in analytical ethanol and shipped to Floragenex Inc. (USA) for genomic DNA extraction, normalisation to 20 ng/μL, paired-end 150bp library preparation and sequencing with an Illumina Novaseq S4 flow cell on the Illumina Novaseq 6000 platform.

Quality of raw data was checked using *FastQC v.0.12.0* (https://www.bioinformatics.babraham.ac.uk/projects/fastqc/). Reads were assigned to individual samples, and duplicates removed, using process_radtags 2.4 and clone_filter 2.4, respectively, within *Stacks* (Catchen et al., 2013). Reads were aligned to the rainbow trout reference genome (Pearse et al., 2019; GenBank Assembly Accession GCA_013265735.3 USDA_OmykA_1.1 2020) using BWA-MEM (Li, 2013). Conversion of sam to bam files was made with *sambamba v.1.0* (Tarasov et al., 2015), and *angsd v.0.941* (Korneliussen et al., 2014) was used to calculate genotype likelihoods. Reads were excluded if they had a mapping quality lower than 30 (-minMApq 30), a flag above 255 (-remove bad 1), did not uniquely map, were not properly paired (-only proper pairs 1); and mapping quality was adjusted for excessive matches (-C 50). Individual positions were only considered if the base quality score was above 20 (-minQ 20), if at least half of all individuals had two reads, the minimum read depth was above 1 × the total number of individuals (min depth of 219) across all individuals, and the maximum read depth was below 2 × mean depth × total number of individuals (maximum depth of 8353). Positions were considered single nucleotide polymorphisms (SNPs) if the SNP *p*-value was below 10e^-6^ and sites had two alleles. SNPs with a rarer allele frequency of less than 5% were further excluded.

### Analyses

All analyses were, unless otherwise indicated, conducted with *R v.4.5.1* (R Core Team, 2025). A principal components analysis (PCA) based on identified SNPs after filtering was conducted using *PCAngsd v.1.2* (Meisner & Albrechtsen, 2018). The same package was used to construct admixture plot showing ancestry proportions for all hatchery and stream populations, after determining the number of genetic clusters that best explain genetic structure, based on ten-fold cross-validation error. A phylogenetic tree was constructed following the neighbour-joining method to visualise relationships between populations using *FigTree v.1.4.4* (http://tree.bio.ed.ac.uk/software/figtree/).

A second genomic PCA was performed in *PCAngsd v.1.2* (Meisner & Albrechtsen, 2018) for the Omy5 region alone, where an inversion has previously been described (Pearse et al., 2019): chromosome Omy5, positions 35,000,000–85,000,000. All individuals were assigned to one of three possible genotypes of the Omy5 inversion, and genotype frequencies (homozygous for the major allele, homozygous for the alternative allele, and heterozygous) were calculated for each population and origin type (*i.e.* hatchery *vs* stream).

Genetic diversity values, *i.e.* nucleotide diversity (*π*) and Watterson’s *θ* (*θ*_w_), were calculated in 100 kb windows across the genome using all sites, variant and invariant, that were present in at least 50% of individuals, had a minimum coverage of 2x per included individual, a maximum coverage of 2 × mean depth (*i.e.* 18.8x) × number of individuals, and passed all other basic filters (*i.e.* mean mapping quality of 30, mean base quality of 20), but without filtering for minor allele frequency. A linear mixed effects model was fitted with the lmer function in the *R*-package, *lme4* (Bates et al., 2015), to test the effect of origin (hatchery *vs* stream) on *π* and *θ*_w_. To control for interpopulation variation in both hatchery and stream groups, specific population was treated as a random factor.

Genetic differentiation (*F*_ST_) was calculated genome-wide using *angsd v.0.941* (Korneliussen et al., 2014) and visualised with a Manhattan plot created in *CMplot v.4.2.0* (Yin et al., 2021). Pairwise comparisons of *F*_ST_ values were made to establish means between hatchery–hatchery, stream–hatchery, and stream–stream populations. Differences in *F*_ST_ between pairwise comparisons were tested with ANOVA followed by Tukey’s HSD *post hoc*.

Finally, we conducted a genome-wide association study (GWAS) to determine positions in the genome that are associated with origin (hatchery or stream). We created a mean genotype file from genotype probabilities and accounted for genetic covariance in a linear mixed model, using the lmm1 command in *GEMMA v.0.98.5* (Zhou & Stephens, 2012) and visualised results with a Manhattan plot created in *CMplot v.4.2.0* (Yin et al., 2021). We determined loci as significant if the *p*-value was below a conservative threshold of 0.00000001 (10^−8^), which is well established for common variants and takes multiple testing into account (Fadista et al., 2016). Loci showing a significant association were manually checked against the rainbow trout genome using the genome browser, *Salmobase* (https://salmobase.org), to identify genes within which the SNPs were found plus adjacent genes.

## Results

### Sampling

In all 14 streams in which allochthonous rainbow trout were sampled, autochthonous brown trout were also caught during electrofishing or angling surveys. In several streams, the target number of 15 sampled rainbow trout was not reached, even after intensive electrofishing (*i.e.* Breg, Lauter and Schiltach; Supplementary material, Fig. S1). In these streams, the population density of brown trout was many times higher than that of rainbow trout. In other streams, comparable numbers of brown and rainbow trout were caught during electrofishing surveys (*e.g.* Kinzig, Wilde Gutach, lower Wutach). In Gutach and upper Wutach, the number of rainbow trout caught via electrofishing was nearly zero, and only repeated fly fishing realised a sufficient sample size.

### Population structure

After filtering, a total of 533,536 SNPs with a mean depth of coverage of ∼18x were identified. The PCA based on SNPs revealed three main clusters on the PC1 and PC2 axes: one containing only the Insenbach/Mühlbach population, another containing Lauter and Lautracher Ach populations, and the third containing the remaining naturalised populations together with hatchery populations (Fig. 1A). Within the large third PCA cluster, individuals from Fischzucht Störk (hatchery) stood slightly apart from the remaining populations. On the PC3 and PC4 axes, the three hatcheries were separated clearly from one another, while the Höllenbach stream population clustered on its own, and there was further fine scale clustering amongst several other populations (Fig. 1B). These population structuring patterns were confirmed by a neighbour-joining phylogenetic tree (Fig. 1C) and the admixture analysis, which found that genetic structure was best explained with 15 genetic clusters (K = 15) (Supplementary materials, Fig. S2).

**Figure 1.**
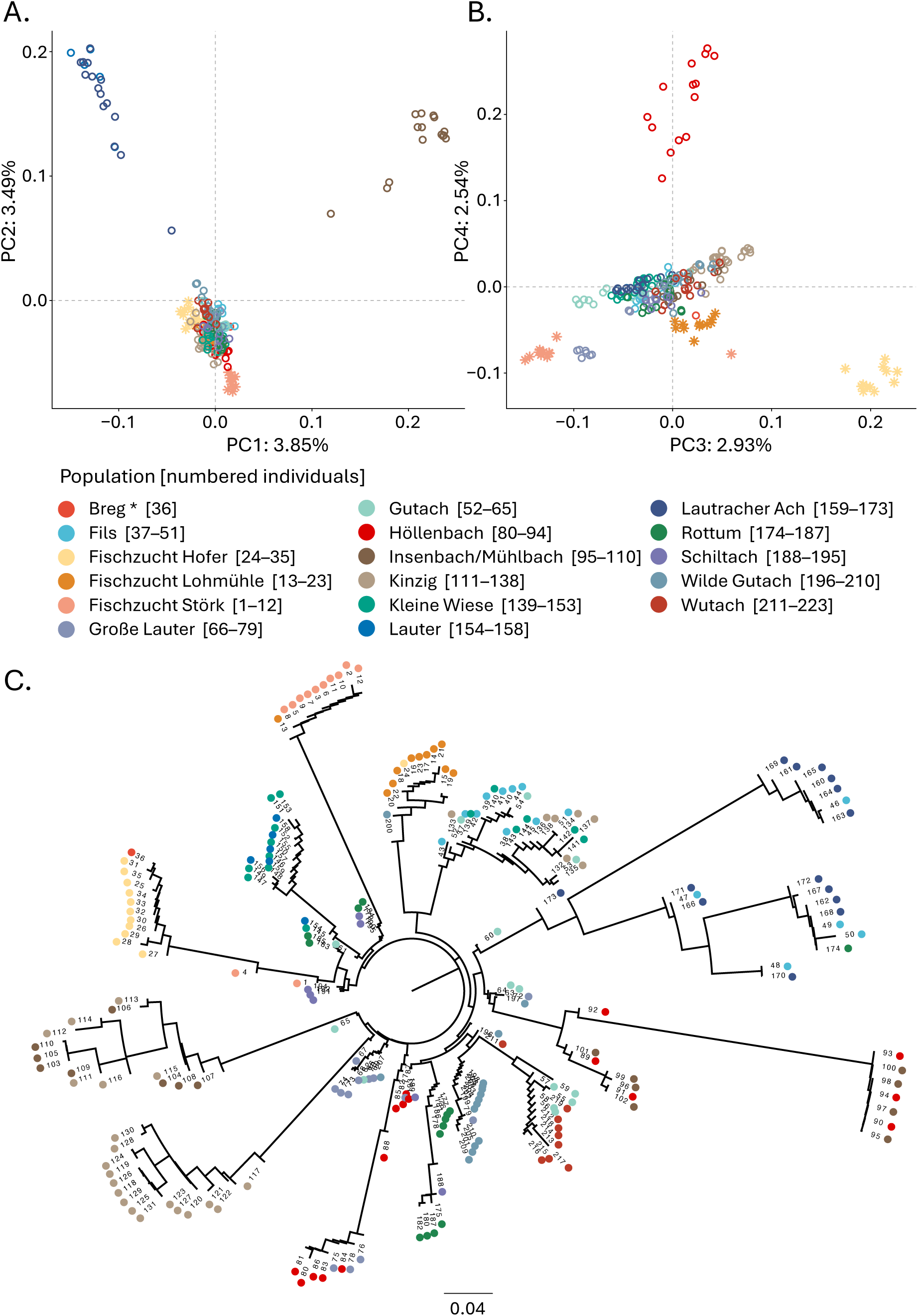
Population structure of 14 stream and 3 hatchery populations of rainbow trout, displayed as (A.) PC1 and PC2 axes, and (B.) PC3 and PC4 axes of a genomic PCA; and (C.) a phylogenetic tree constructed by neighbour-joining method.

Regarding the Omy5 inversion, which was chosen for particular consideration because of its association with life history variants, three distinct clusters showed clearly on the PC1 axis, explaining a large proportion (33.5%) of genetic variation in this region (Fig. 2A). The pattern corresponds to three genotype variants, which were assumed to represent the three inversion types: homozygous for the major allele, homozygous for the alternative allele, and heterozygous. All three genotypes were present in both hatchery and stream populations, without any clear difference in inversion frequencies between hatchery and stream populations (Fig. 2B).

**Figure 2.**
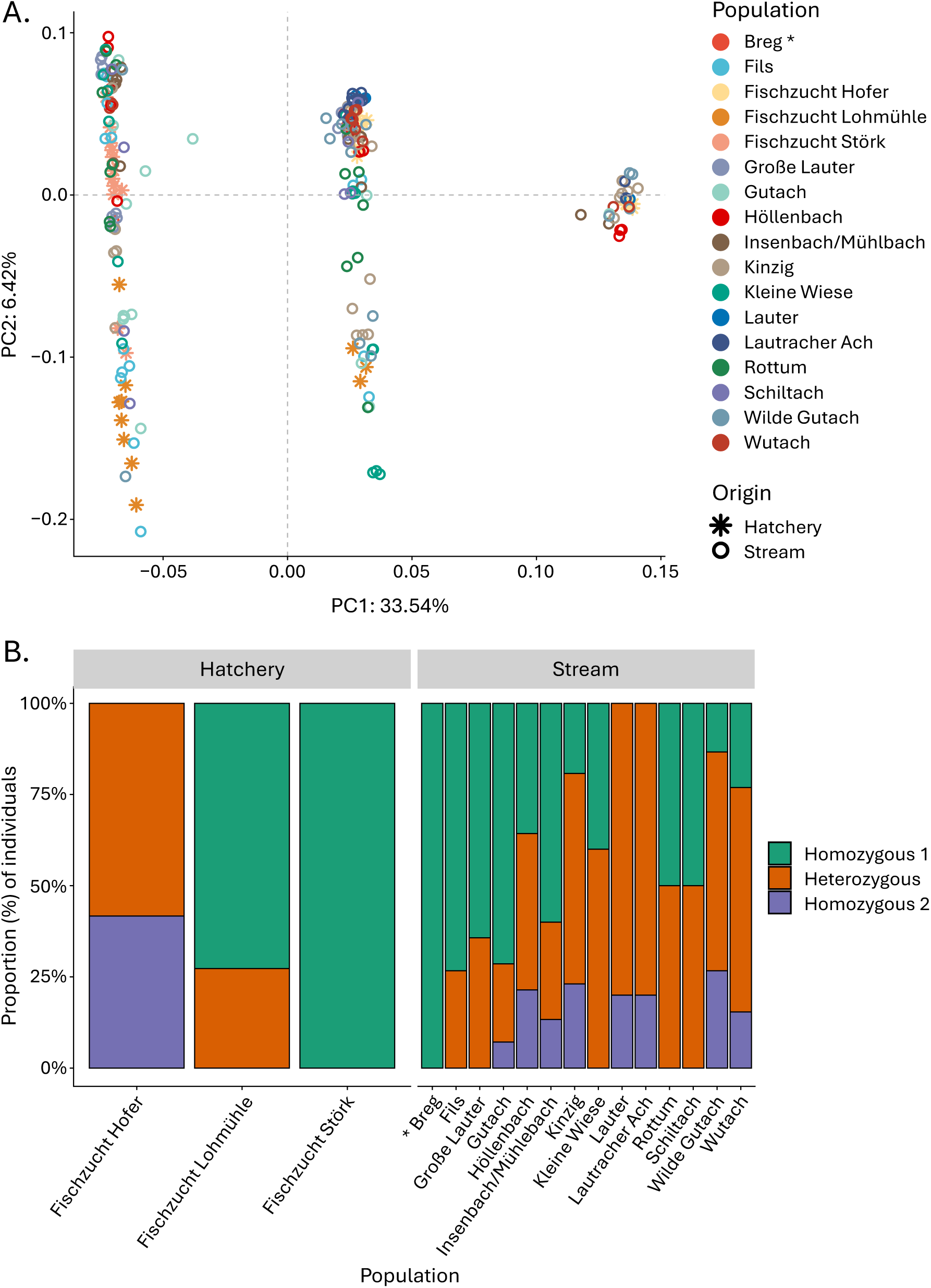
Variation in the Omy5 chromosome across 14 stream and 3 hatchery populations of rainbow trout: (A.) clustering in PCA; (B.) proportion of individuals by population homozygous with the major allele, alternate allele, or heterozygous. * Nb. Breg is represented by a single individual.

### Genetic diversity

Although we found variation between individual populations, there was no significant difference in genetic diversity across the genome, generally, between hatchery and stream trout: the effect of origin (stream) on *π* was β = 2.43e^−4^, 95% CI [−5.4e^−5^, 5.4e^−4^], t_297695_ = 1.6, *p* = 0.109; Std. β = 0.13, 95% CI [−0.03, 0.29]; the effect of origin (stream) on *θ*_w_ was β = 2.66e^−4^, 95% CI [−1.01e^−4^, 6.34e^−4^], t_297695_ = 1.42, *p* = 0.156; Std. β = 0.18, 95% CI [−0.07, 0.42] (Supplementary material, Figs. S3 and S4).

### Genetic differentiation

Genetic differentiation was pronounced for all population comparisons (Fig.3; Supplementary Material, Table S1), with the greatest distinction between hatchery populations (mean *F*_ST_ ± s.d. = 0.1965 ± 0.0382), followed by stream–hatchery comparisons (mean *F*_ST_ ± s.d. = 0.1715 ± 0.0437), and the least differentiation between stream populations (mean *F*_ST_ ± s.d. = 0.1377 ± 0.049). There was a significant effect of comparison on weighted *F*_ST_ (*F*_2,116_ = 8.11, *p* < 0.001), with the difference between the stream–stream comparison significantly lower than the stream–hatchery comparison (difference = −0.0337, 95% CI [−0.0557, −0.0117], *p* = 0.001) (Fig. 3). However, there was no strong signal of differentiation across the Omy5 inversion (Fig. 4A), as suggested by Omy5 inversion frequencies in individual hatchery and stream populations. We detected widespread genetic differentiation across the genome between hatchery and stream populations with genomic regions on most chromosomes showing differentiation above the 99^th^ percentile of the *F*_ST_ distribution (*F*_ST_ > 0.259). Between individual populations, Insenbach/Mühlbach was most differentiated from the hatcheries (from Fischzucht Hofer, *F*_ST_ = 0.2804, from Fischzucht Lohmühle, *F*_ST_ = 0.2271, and from Fischzucht Störk, *F*_ST_ = 0.2543), followed by Lauter (from Fischzucht Hofer, *F*_ST_ = 0.2415, from Fischzucht Lohmühle, *F*_ST_ = 0.1916, and from Fischzucht Störk, *F*_ST_ = 0.2308), and Lautracher Ach (from Fischzucht Hofer, *F*_ST_ = 0.2170, from Fischzucht Lohmühle, *F*_ST_ = 0.1725, and from Fischzucht Störk, *F*_ST_ = 0.2101). *F*_ST_ values between Insenbach/Mühlbach and both Lauter and Lautracher Ach were larger than average (*F*_ST_ = 0.2772 and 0.2504, respectively). There was, however, little difference between Lauter and Lautracher Ach (*F*_ST_ = 0.0053).

**Figure 3.**
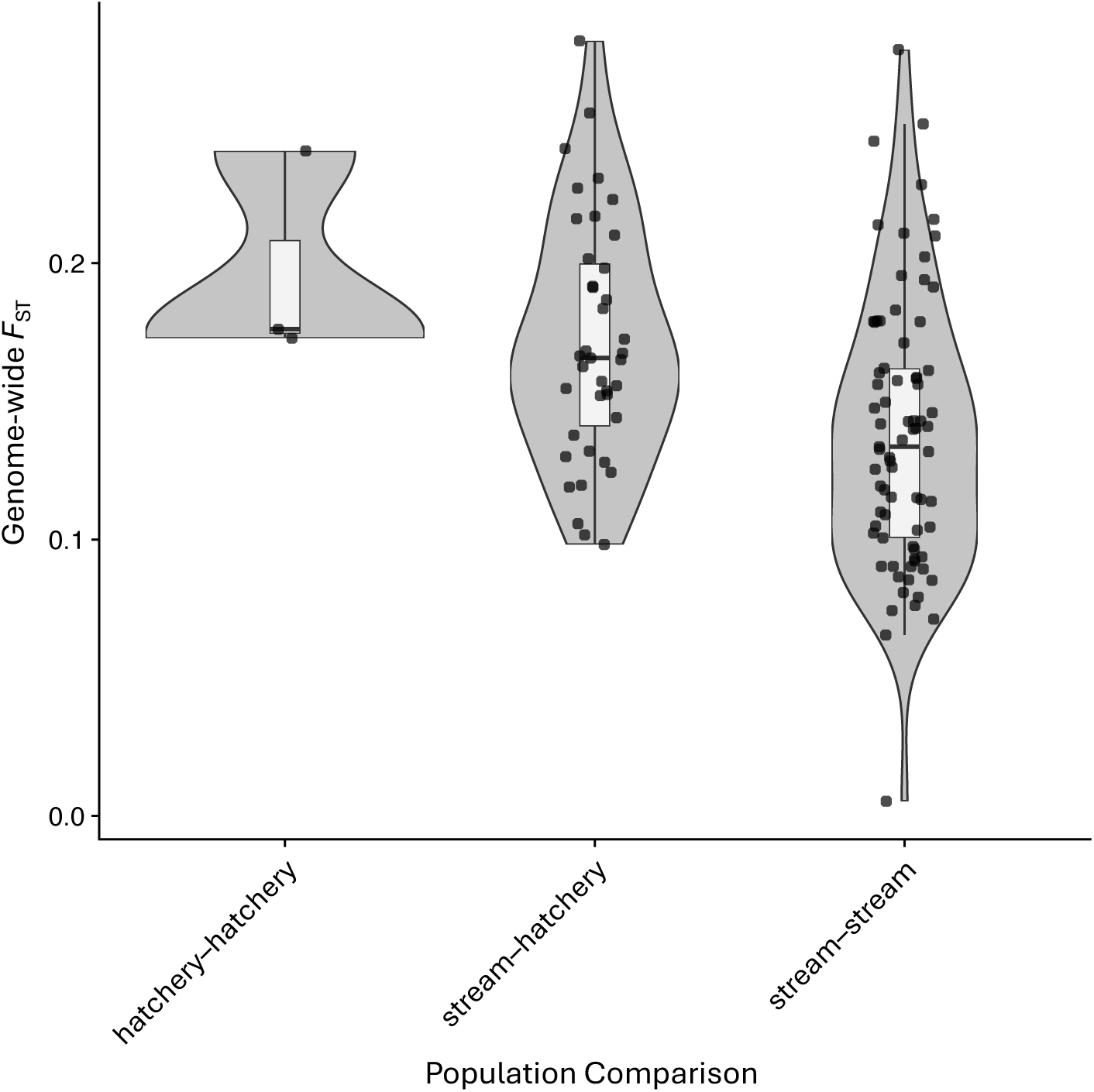
Effect of comparison on weighted *F*_ST_ across the genome. Comparisons were made between 3 hatchery populations, 14 stream populations and hatchery and stream populations of rainbow trout.

**Figure 4.**
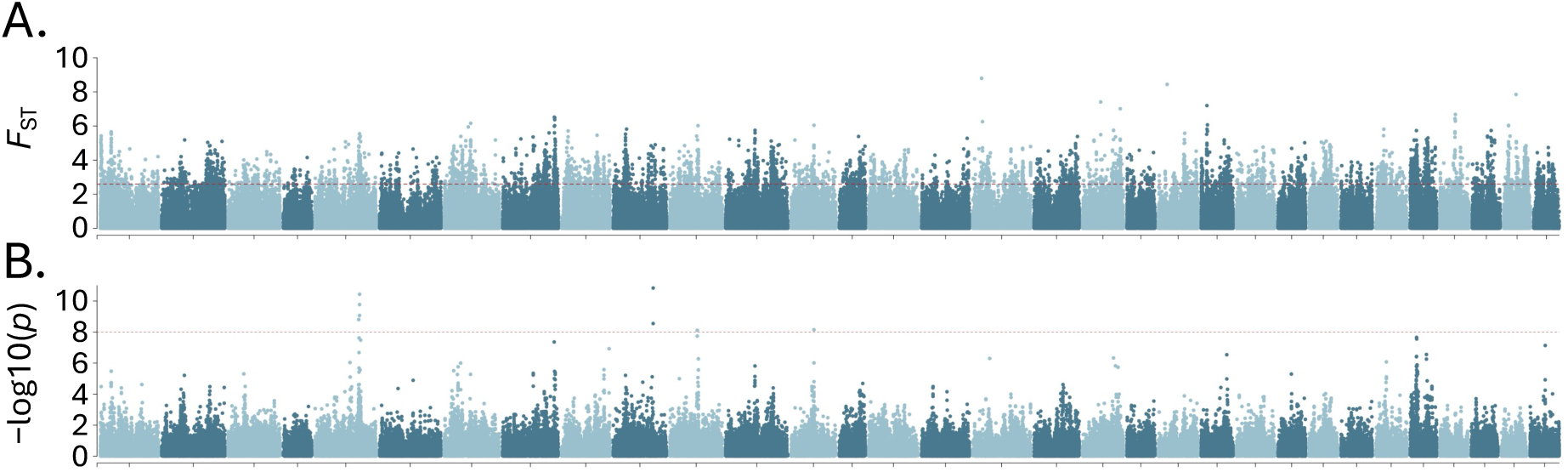
Manhattan plots showing (A.) genetic differentiation (*F*_ST_) across the genome, and (B.) values of −log(10) of *p* from a genome-wide association study of 3 hatchery and 14 stream populations of rainbow trout. Chromosomes are coloured in alternating order for visual clarity. The dashed red line in (A.) shows the 99^th^ percentile of the genome *F*_ST_ distribution (*F*_ST_ = 0.259) and the dashed red line in (B.) shows the *p*-value threshold for significant genome-wide estimation (*p* = 0.00000001).

### Genome-wide association with origin

The GWAS identified nine SNPs significantly associated with origin (hatchery *vs* stream), four of which were found within a narrow peak inside the Omy5 inversion. The other SNPs were located on chromosomes Omy10, Omy11 and Omy13. These origin-associated regions also showed increased genetic differentiation between hatchery and stream populations. Significant GWAS SNPs were located within or close to genes encoding for proteins involved in amino acid and protein transport; phosphatase activity; development and proliferation of T-cells; neural and cardiac development; and cell growth, proliferation, homeostasis and apoptosis (Fig. 4B; Supplementary Materials: Table S2).

## Discussion

Population structuring amongst the rainbow trout populations in the examined streams of Baden-Württemberg is generally consistent with the assumption that these populations reproduce naturally and are self-sustaining: although most populations appear to be related, structuring is present throughout the region. Most of the stream populations showed ancestry from all three hatchery populations considered here, suggesting that these hatcheries were indeed source populations for most of the streams, with only Lauter and Lautracher Ach as exceptions. This is further supported by the lower genetic differentiation between stream populations compared to the hatchery populations, which might suggest a mixed hatchery origin of stream populations, potentially through genetic mixing caused by multiple stocking events or post-colonisation gene flow between streams. A reduction in genetic diversity (as opposed to differentiation) is a commonly recognised signature of founder events, in which only a portion of an original source population’s genetic diversity is carried to a novel environment; however, the loss can be mitigated by the number of colonisation events and sources (Barton & Charlesworth, 1984; Roman, 2006). In comparing the stream rainbow trout to the hatchery populations, no evidence was found of reduced genetic diversity, further suggesting that multiple sources contributed to their naturalisation (Lázari et al., 2024), which is reflected in the substantial admixture found in the stream populations.

Three stream populations stood out as particularly distinct both from other stream populations and the hatcheries. However, the mechanisms of differentiation in the populations at Insenbach/Mühlbach, Lauter and Lautracher Ach can only be subject to speculation. One possibility is that these populations have arisen from other breeding lineages of different geographic origin or hatcheries not considered here – potentially additional source lineages, as indications are that the Lauter and Lautracher Ach populations share a common ancestry. It may be that differing local environmental conditions in individual streams may present some unidentified selection pressure that promotes adaptation, which can occur over relatively few generations (Schluter, 2009), and has been documented after historical stocking events in salmonids (e.g. Crotti et al., 2021; Koene et al., 2019; Vuorinen, 1991). It is also possible that the differentiation of these three populations results from founder effects of a small historical stocking input followed by genetic drift (Hauser et al., 1995; Weeder et al., 2005), as has been described in other rainbow trout naturalisation events (Colihueque et al., 2019), although the genetic diversity of the stream populations would appear to argue against this (Austerlitz et al., 1997). Comparisons to other putative source populations would be required to distinguish between scenarios; however, we are unaware of any other potential source populations, and so were unable to include these in the study.

Three genotypes of the Omy5 inversion associated with life history were present amongst the stream populations, potentially corresponding to the homozygous anadromous, homozygous resident and heterozygous genotypes described in the literature (Goetz et al., 2024; Pearse et al., 2014). However, without sequence material from an individual with a known migratory phenotype, it cannot be determined which phenotypes associate with genotypes uncovered here. Nonetheless, we found no strong *F*_ST_ signal within the Omy5 inversion. The frequencies of genotypes were not consistent across either hatchery or stream populations, and it is therefore not possible to evaluate the role that the inversion plays in the naturalisation of rainbow trout in wild streams.

On the other hand, of the nine positions identified by the GWAS to be associated with naturalisation, four were found within the Omy5 inversion. These positions were found within, or adjacent to, genes associated with immunity and growth and development, which are presumably under different selection pressures in the wild compared to hatchery environments, and may be related to phenotypes conducive to naturalisation (Kanno et al., 2017; Lázari et al., 2024). Consistent with our hypothesis that genetics play a role in determining whether rainbow trout can naturalise in wild ecosystems, we found that certain genetic variants are more likely than others to become established in the wild, which points to potential drivers of adaptation. It has been stated that the successful establishment of self-sustaining rainbow trout populations is driven primarily by genetic factors (Koutsikos et al., 2019), but that environmental conditions play important roles, too (Fausch et al., 2001); we cannot rule out the possible importance of environmental variables. Furthermore, we found no evidence that the descendants of any one of the three hatchery populations had become more prevalent than the others in the wild, thereby posing a particular risk of spreading an invasive species. However, we cannot discount the possibility of a prolific source population, which has simply escaped our notice.

Our finding that the genetic profiles of all sampled hatchery stock are broadly represented in the rainbow trout of the different streams, has important implications for the management of local trout waters, as the source of the stocking material appears not to be as important as hypothesised for the future establishment of rainbow trout stock. Because different stocked trout had come from hatchery stocks intensively selected for aquaculture optimisations, we hypothesised that the longer the selection history and/or the more artificial the rearing environment, the smaller the chance of establishment in the wild would be (Petersson et al., 1996; Lorenzen et al., 2012). However, our data did not support this hypothesis, and current levels of domestication or husbandry practice, at least, do not appear to prevent naturalisation probabilities: regardless of stocking source or time span of domestication, all stocked rainbow trout appear to have the potential to establish in nature. However, we must acknowledge that we sampled only three different farms with distinct stock characteristics, and that further investigation is needed for a more in-depth understanding of the properties important for naturalisation.

Furthermore, the intensity of postulated competition with native brown trout (Stanković et al., 2016) has to be questioned. Despite more than 100 years of intensive rainbow trout stocking in southwestern Germany, extensive research was required to determine the areas in which reproducing populations occur. Although many, possibly even all, trout streams in Baden-Württemberg had been increasingly stocked with rainbow trout over the past century, in only a few places have rainbow trout been able to establish self-sustaining populations. Additionally, we found brown trout still present in all streams, where they co-exist with rainbow trout. Rainbow trout were never dominant; indeed, often it was the brown trout populations that predominated. In many streams, rainbow trout establishment has occurred at very low levels, and in some instances with no apparent signs of local population expansion (*i.e.* increase in density or range) in more than 20 years, nor with signs of negative effects on brown trout populations (Baer & Brinker, 2010). Therefore, the widely postulated phenomenon of competition, based mainly on studies from countries where both species are allochthonous (Hayes, 1987), appears to exist only to a limited extent, at most, in the investigated ecoregion. However, some recent facilitation dynamic is evident, as documentation of self-sustaining rainbow trout populations is limited to recent decades, despite more than a century of intense stocking. This study indicates that the propagule pressure framework, which relates invader establishment success to the density of introduced individuals, the number of introduction events, and the relative frequency of introductions (Simberloff, 2009), does not adequately explain the patterns observed in this species and ecoregion. Instead, the invasion potential of rainbow trout has likely increased recently through, for example, the emergence of additional invasive species and/or the effects of climate change. Further research is therefore required to identify the conditions under which rainbow trout can successfully naturalise, and to quantify any resulting impacts on local brown trout populations.

In summary, we found clear population structuring of rainbow trout populations in the streams of Baden-Württemberg, which provided evidence of natural reproduction over several generations. Multiple genetic origins of stream populations were inferred: the three hatcheries under consideration here, and possibly others; and the origins were likely mixed via pre- or post-colonisation gene flow. However, despite multiple origins, there was no significant difference in genetic diversity between stream and hatchery populations, but there were specific positions in the genome associated with naturalisation. Fisheries management tasked with sustaining native salmonids affected by allochthonous rainbow trout should benefit from more detailed knowledge concerning rainbow trout population structure and origins.

## Acknowledgments

We are grateful to Laura Epp for supporting the project to come to life in its early stages. We also thank the officials of Baden-Württemberg Fisheries Authority and fishing wardens for their frequent and helpful information regarding streams with self-sustaining rainbow trout populations. Furthermore, we offer very special thanks to the team at FFS Langenargen, particularly Phillipp Tölle, for their support and help in catching trout and sampling. We express our gratitude to the local angling clubs and tenantry for their participation in the project: *ASV Elsenztal Steinsfurt 1968 e.V.*, *ASV Freiburg e.V.*, *ASV Gutach e.V.*, *AV Simonswäldertal e.V.*, *ASV Tennenbronn, AV Wolterdingen e.V.*, *Fliegenfischervereinigung 1966 Wutach-Flühen e.V.*, *FV Ochsenhausen e.V.*, Hannes Junger, Franz Mühlhans, Dieter Niemietz, and Karl Skrodski. Finally, we thank Stephan Hofer (Fischzucht Hofer), Joachim Schinldler (Fischzucht Lohmühle) and Peter Störk (Fischzucht Störk) for providing hatchery samples.

## Data availability statement

All environmental and ecological data are included within either this article or its supplementary materials. Sequence reads will, upon publication, be archived at the NCBI Sequence Read Archive.

## Funding

This project was funded by the Fischereiabgabe Baden-Württemberg.

### Ethics statement

All fish were caught by licenced personnel under permission of the local fisheries administration (Regierungspräsidium Karlsruhe, Freiburg, Tübingen) according to the German Animal Protection Law (§ 4). Captured fish were killed according to the ordinance on slaughter and killing of animals (Tierschutzschlachtverordnung § 13).

### Author contributions (CRediT)

Conceptualisation: AB Methodology: PB, AB Validation: PB, AB

Formal analysis: JPK, AJ, DF, PV, AB Investigation: PB, JB

Data curation: PB, JB, DF, PV Writing – original draft: JPK, JB

Writing – review & editing: JPK, AJ, PB, JB, DF, PV, AB Visualisation: JPK, AJ

Supervision: AB

Project administration: AB Funding acquisition: JB, AB

## References

Arlinghaus, R., Mehner, T., & Cowx I.G. (2002). Reconciling traditional inland fisheries management and sustainability in industrialised countries, with an emphasis on Europe. Fish and Fisheries, 3(4), 261–316. 10.1046/j.1467-2979.2002.00102.x

Austerlitz, F., Jung-Muller, B., Godelle, B., & Gouyon, P.-H. (1997). Evolution of coalescence times, genetic diversity and structure during colonisation. Theoretical Population Biology, 51(2), 148–164. 10.1006/tpbi.1997.1302

Baer, J. & Brinker, A. (2010). The response of a brown trout stocks and perception of anglers to cessation of brown trout stocking. Fisheries Management and Ecology, 17(2), 157–164. 10.1111/j.1365-2400.2009.00713.x

Barton, N.H. & Charlesworth, B. (1984). Genetic revolutions, founder effects, and speciation. Annual Review of Ecology and Systematics, 15, 133–164. 10.1146/annurev.es.15.110184.001025

Bates, D., Mächler, M., Bolker, B., & Walker, S. (2015). Fitting linear mixed-effects models using lme4. Journal of Statistical Software, 67, 1–48. 10.18637/jss.v067.i01

Burkhardt-Holm, P., Peter, A., & Segner, A.H. (2002). Decline of fish catch in Switzerland. Aquatic Sciences, 64, 36–54. 10.1007/s00027-002-8053-1

Catchen, J., Hohenlohe, P.A., Bassham, S., Amores, A., & Cresko, W.A. (2013). Stacks: an analysis tool set for population genomics. Molecular Ecology, 22(11), 3124–3140. 10.1111/mec.12354

Colihueque, N., Estay, F.J., Crespo, J.E., Arriigada, A., Baessolo, L., Canales-Aguire, C.B., Marín, J., & Carrasco, R. (2019). Genetic differentiation and origin of naturalised rainbow trout populations from southern Chile, revealed by the mtDNA control region marker. Frontiers in Genetics, 10, 1212. 10.3389/fgene.2019.01212

Crawford, S.S. & Muir, A.M. (2008). Global introductions of salmon and trout in the genus *Oncorhynchus*: 1870–2007. Revies in Fish Biology and Fisheries, 18, 313–344. 10.1007/s11160-007-9079-1

Crotti, M., Yohannes, E., Winfield, I.J., Lyle, A.A., Adams, C.E., & Elmer, K.R. (2021). Rapid adaptation through genomic and epigenomic responses following translocations in an endangered salmonid. Evolutionary Applications, 14(10), 2470–2489. 10.1111/eve.13267

Elmer, K.R. & Meyer, A. (2011). Adaptation in the age of ecological genomics: insights from parallelism and convergence. Trends in Ecology & Evolution, 26(6), 298 – 306. 10.1016/j.tree.2011.02.008

Fadista, J, Manning, A.K., Florez, J.C., & Groop, L. (2016). The (in)famous GWAS *P*-value threshold revisted and updated for low-frequency variants. European Journal of Human Genetics, 24, 1202 – 1205. 10.1038/ejhg.2015.269

Fausch, K.D. (2007). Introduction, establishment and effects of non-native salmonids: considering the risk of rainbow trout invasion in the United Kingdom. Journal of Fish Biology, 71(sd) 1–32. 10.1111/j.1095-8649.2007.01682.x

Fausch, K.D., Taniguchi, Y., Nakano, S., Grossman, G.D., & Townsend, C.R. (2001). Flood disturbance regimes influence rainbow trout invasion success among five Holarctic regions. Ecological Applications, 11(5), 1438–1455. 10.1890/1051-0761(2001)011[1438:FDRIRT]2.0CO;2

Gatz, A.J., Sale, M.J., & Loar, J.M. (1987). Habitat shifts in rainbow trout: competitive influences of brown trout. Oecologia, 74, 7–19. 10.1007/BF00377339

Goetz, L.C., Nuetzel, H., Vendrami, D.L.J., Beulke, A.K., Anderson, E.C., Garza, J.C., & Pearse, D.E. (2024). Genetic parentage reveals the (un)natural history of Central Valley hatchery steelhead. Evolutionary Applications, 17, e13681. 10.1111/eva.13681

Halverson, A. (2010). An entirely synthetic fish: How rainbow trout beguiled America and overran the world. New Haven & London: Yale University Press. 10.2307/j.ctt1nq8bk

Hauser, L., Carvalho, G.R., & Pitcher, T.J. (1995). Morphological and genetic differentiation of the African clupeid *Limnothrissa miodon* 34 years after its introduction to Lake Kivu. Journal of Fish Biology, 47(sA), 127–144. 10.1111/j.1095-8649.1995.tb06049.x

Hayes, J.W. (1987). Competition for spawning space between brown (*Salmo trutta*) and rainbow trout (*S. gairdneri*) in a lake inlet tributary, New Zealand. Canadian Journal of Fisheries and Aquatic Sciences, 44(1), 40–47. 10.1139/f87-005

Kanno, Y., Kulp, M.A., Moore, S.E., & Grossman, G.D. (2017). Native brook trout and invasive rainbow trout respond differently to seasonal weather variation: Spawning time matters. Freshwater Biology, 62(5), 868–879. 10.1111/fwb.12906

Koene, J.P., Crotti, M., Elmer, K.R., & Adams, C.E. (2019). Differential selection pressures result in a rapid divergence of donor and refuge populations of a high conservation value freshwater fish *Coregonus lavaretus* (L.). Evolutionary Ecology, 33, 533–548. 10.1007/s10682-019-09995-y

Korneliussen, T.S., Albrechtsen, A., & Nielsen, R., (2014). ANGSD: Analysis of next generation sequencing data. BMC Bioinformatics, 15, 356. 10.1186/s12859-014-0356-4

Koutsikos, N., Vardakas, L., Zogaris, S., Perdikaris, C., Kalantzi, O.-I., & Economou, A.N. (2019). Does rainbow trout justify its high rank among alien invasive species? Insights from a nationwide survey in Greece. Aquatic Conservation: Marine and Freshwater Ecosystems, 29(3), 409–423. 10.1002/aqc.3025

Lázari, C., Riva-Rossi, C., Ciancio, J., Pascual, M., Clement, A.J., Pearse, D.E., & Garza, J.C. (2024). Ancestry and genetic structure of resident and anadromous rainbow trout (*Oncorhynchus*) in Argentina. Journal of Fish Biology, 104(6), 1972–1989. 10.1111/jfb.15722

Li, H. (2013). Aligning sequence reads, clone sequences and assembly contigs with BWA-MEM. arXiv:1303.3997 [*q-bio,GN*]. 10.48550/arXiv.1303.3997

Lorenzen, K., Beveridge, M.C.M., & Mangel, M. (2012). Cultured fish: integrative biology and management of domestication and interactions with wild fish. Biological Reviews, 87(3), 639–660. 10.1111/j.1469-18X.2011.00215.x

Lowe, S., Browne, M., Boudjelas, S. & De Porter, M. (2000). 100 of the world’s worst invasive alien species: A selection from the Global Invasive Species Database. *ISSG / SSE / IUCN*. Available at https://www.issg.org/booklet.pdf

McGlade, C.L.O., Dickey, J.W.E., Kennedy, R., Donnelly, S., Nelson, C.-A., Dick, J.T.A., & Arnott, G. (2022). Behavioural traits of rainbow trout and brown trout may help explain their differing invasion success and impacts. Scientific Reports, 12, 1757. 10.1038/s41598-022-05484-5

Meisner, J. & Albrechtsen, A. (2018). Inferring population structure and admixture proportions in low-depth NGS data. Genetics, 210(2), 719–731. 10.1534/genetics.118.301336

Mueller, M., Egg, L., Ruff, T., Haas, A., Schubert, M., & Gum, B. (2025). Evidence of natural reproduction of North American rainbow trout (*Oncorhynchus mykiss*) in three Alpine rivers in Bavaria, Germany. Fish Management and Ecology, 32(2), e12765. 10.1111/fme.12765

Pearse, D.E., Barson, N.J., Nome, T., Gao, G., Campbell, M.A., Abadia-Carduso, A., et al. (2019). Sex-dependent dominance maintains migration supergene in rainbow trout. Nature Ecology & Evolution, 3, 1731–1742. 10.1038/s41559-019-1044-6

Pearse, D.E., Miller, M.R., Abadía-Cardoso, A., & Garza, J.C. (2014). Rapid parallel evolution of standing variation in a single, complex, genomic region is associated with life history in steelhead/rainbow trout. Proceedings of the Royal Society B, 281(1783), 20140012. rspb.2014.0012

Petersson, E., Jauvri, T., Steffner, N.G., & Ragnarsson, B. (1996). The effect of domestication on some life history traits of sea trout and Atlantic salmon. Journal of Fish Biology, 48(4), 776–791. 10.1111/j.1095-8649.1996.tb01471.x

R Core Team. R: A language and environment for statistical computing. R Foundation for Statistical Computing, Vienna, Austria. 2025. https://www.R-project.org/

Roman, J. (2006). Diluting the founder effect: Cryptic invasions expand a marine invader’s range. Proceedings of the Royal Society B, 273(1600), 2453–2459. 10.1098/rspb.2006.3597

Ros, A., Schmidt-Posthaus, H., & Brinker, A. (2022). Mitigating human impacts including climate change on proliferative kidney disease in salmonids of running waters. Journal of Fish Diseases, 45, 497–521. 10.1111/jfd.13585

Rubin, A., Bailey, C., Strepparava, N., Wahli, T., Segner, H., & Rubin, J.-F. (2022). Reliable field assessment of proliferative kidney disease in wild brown trout, *Salmo trutta*, populations: When is the optimal sampling period? Pathogens, 11(6), 681. 10.3390/pathogens11060681

Schluter, D. (2009). Evidence for ecological speciation and its alternative. Science, 323(5915), 737–741. 10.1126/science.1160006

Simberloff, D. (2009). The role of propagule pressure in biological invasions. Annual Review of Ecology, Evolution, and Systematics, 40, 81–102. 10.1146/annurev.ecolsys.110308.120304

Stanković, D., Crivelli, A.J., & Snoj, A. (2015). Rainbow trout in Europe: introduction, naturalisation, and impacts. Reviews in Fisheries Science & Aquaculture, 23(1), 39–71. 10.1080/23308249.2015.1024825

Stanković, D., Stephens, M.R., & Snoj, A. (2016). Origin and introduction history of self-sustaining rainbow trout populations in Europe as inferred from mitochondrial DNA and a Y-linked marker. Hydrobiologia, 770, 129–144. 10.1007/s10750-015-2577-6

Tarasov, A., Vilella A.J., Cuppen, E., Mijman, I.J., & Prins, P. (2015). Sambamba: fast processing of NGS alignment formats. Bioinformatics, 31(12), 2032–2034. 10.1093/bioinformatics/btv098

Vuorinen, J., Næsje, T.F., & Sandlund, O.T. (1991). Genetic changes in a vendace Coregonus Albula (L.) population 92 years after introduction. Journal of Fish Biology, 39(sA), 193–201. 10.1111/j.1095-8649.1991.tb05083.x

Weber, G.M., Martin, K.E., Palti, Y., Liu, S., Beach, J.N., & Birkett, J.E. (2023). Effects of fertilizing eggs from a summer-spawning line with cryopreserved milt from a winter-spawning line on spawning date and egg production traits in rainbow trout. Aquaculture Reports, 29, 101495. 10.1016/j.aqrep.2023.101495

Weeder, J.A., Marshall, A.R., & Epifanio, J.M. (2005). An assessment of population genetic variation in chinook salmon from seven Michigan rivers 30 years after introduction. North American Journal of Fisheries Management, 25(3), 861–875. 10.1577/M03-227.1

Yin, L., Zhang, H., Tang, Z., Xu, J., Yin, D., Zhang, Z. et al. (2021). rMVP: a memory-efficient, visualization-enhanced, and parallel-accelerated tool for genome-wide association study. *Genomics*, Proteomics & Bioinformatics, 19(4), 619–628. 10.1016/j.gpb.2020.10.007

Young, K.A., Dunham, J.B., Stephenson, J.F., Terreau, A., Thailly, A.F., Gajardo, G., & Garcia de Leaniz, C. (2010). A trial of two trouts: comparing the impacts of rainbow and brown trout on a native galaxiid. Animal Conservation, 13(4), 399–410. 10.1111/j.1469-1795.2010.00354.x

Zhou, X. & Stephens, M. (2012). Genome-wide efficient mixed-model analysis for association studies. Nature Genetics, 44, 821–824. 10.1038/ng.2310

