## Supplementary material for "Multiple genetic origins of non-native, self-sustaining rainbow trout *Oncorhynchus mykiss* in streams in Baden-Württemberg, Germany"

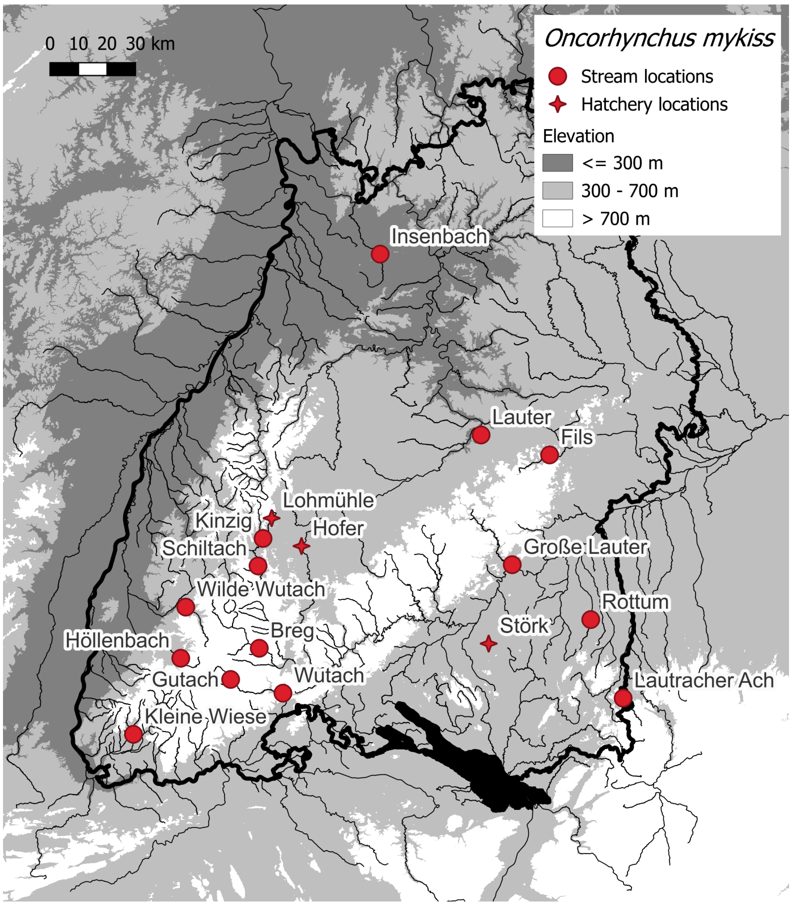


| Stream populations | *n* |
|---|---|
| Breg | 1 |
| Fils | 14 |
| Große Lauter | 12 |
| Gutach | 14 |
| Höllenbach | 14 |
| Insenbach/Mühlbach | 15 |
| Kinzig | 25 |
| Kleine Wiese | 14 |
| Lauter | 5 |
| Lautracher Ach | 15 |
| Rottum | 14 |
| Schiltach | 8 |
| Wilde Gutach | 15 |
| Wutach | 13 |

| Hatchery populations: | *n* |
|---|---|
| Fischzucht Hofer | 12 |
| Fischzucht Lohmühle | 11 |
| Fischzucht Störk | 12 |

**Figure S1.** Sampling locations of 14 stream and 3 hatchery populations of rainbow trout (total *n* = 219) in Baden-Württemberg, Germany.

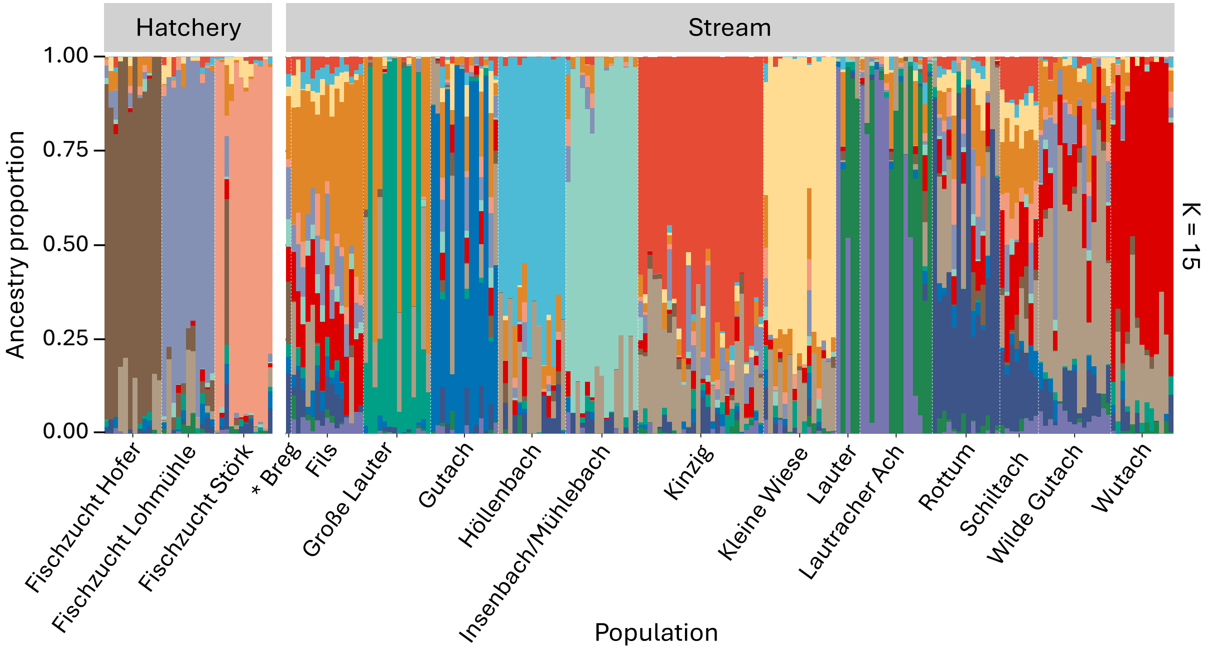


**Figure S2.** Admixture analysis across 3 hatchery and 14 wild stream populations of rainbow trout. Genetic structure was best explained with 15 genetic clusters (K = 15), based on ten-fold cross-validation error.

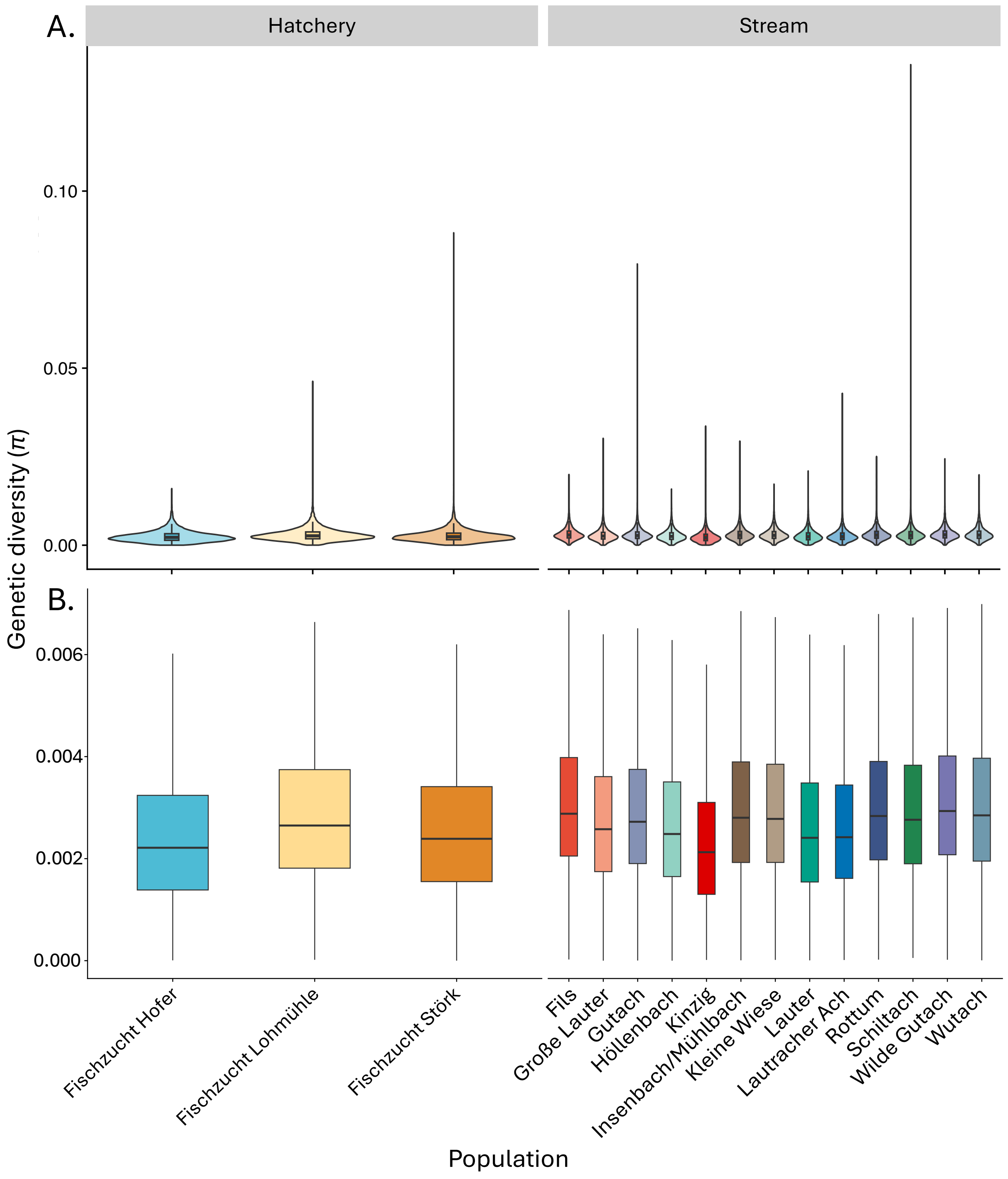


**Figure S3.** Genetic diversity measured as $\pi$ in 100 kb windows across the genome of 3 hatchery and 13 stream populations of rainbow trout. One stream population (Breg) was omitted because it was represented by a single individual. Outliers are included in (A), and excluded in (B).

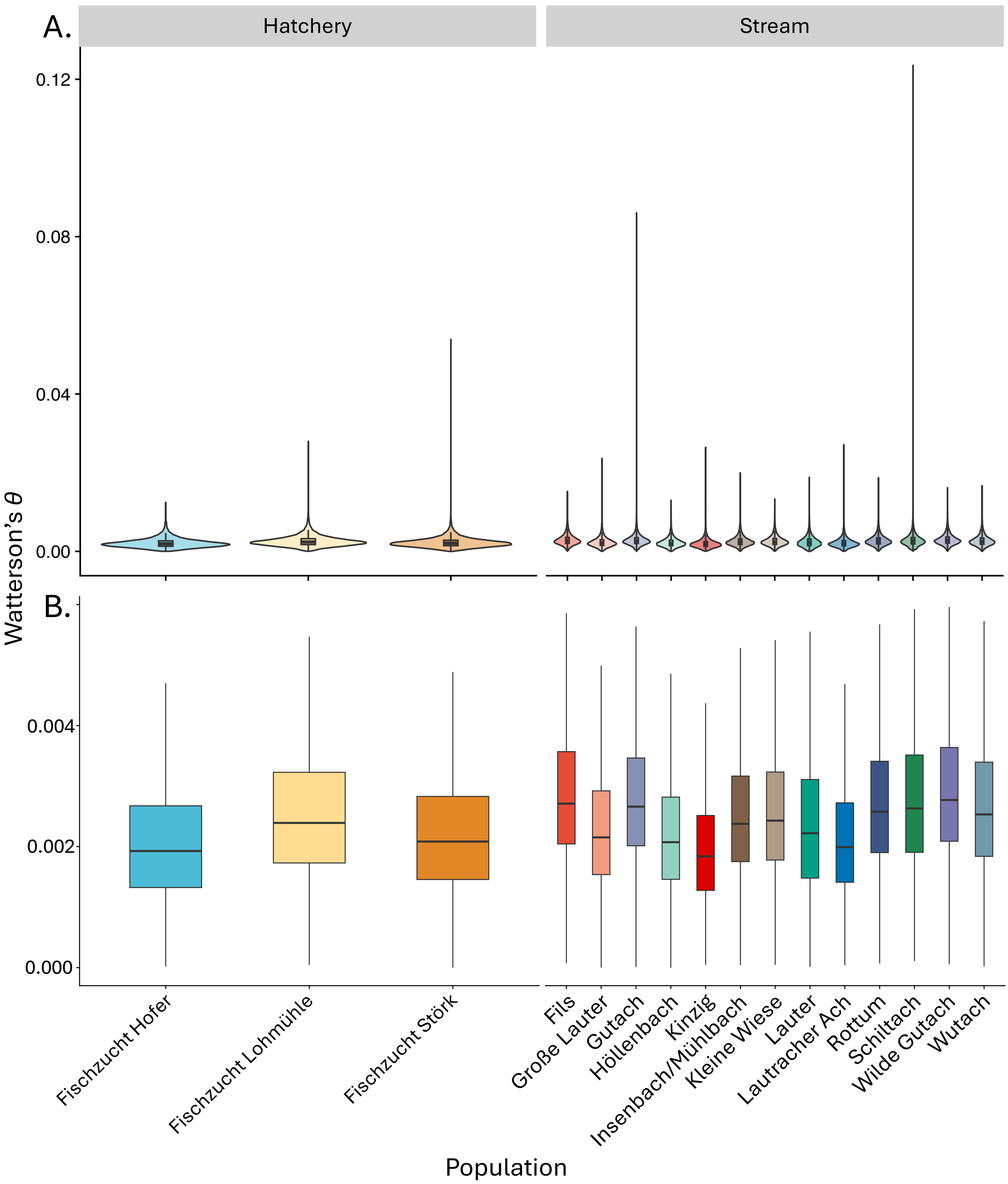


**Figure S4.** Genetic diversity measured as Watterson's $\theta$ in 100 kb windows across the genome of 3 hatchery and 13 stream populations of rainbow trout. One stream population (Breg) was omitted because it was represented by a single individual. Outliers are included in (A), and excluded in (B).

**Table S1.** Weighted FST between comparisons of 3 hatchery and 14 stream populations of rainbow trout.

| Comparison | Pop1 | Pop2 | Weighted_Fst |
|---|---|---|---|
| stream_hatchery | fils | fischzucht_hofer | 0.167391 |
| stream_hatchery | fils | fischzucht_lohmuehle | 0.098207 |
| stream_hatchery | fils | fischzucht_stoerk | 0.131987 |
| stream_stream | fils | grosse | 0.10232 |
| stream_stream | fils | gutach | 0.093727 |
| stream_stream | fils | hoellenbach | 0.128403 |
| stream_stream | fils | insenbach | 0.178828 |
| stream_stream | fils | kinzig | 0.085261 |
| stream_stream | fils | kleine_Wiese | 0.086521 |
| stream_stream | fils | lauter | 0.147564 |
| stream_stream | fils | lautracher_Ach | 0.132678 |
| stream_stream | fils | rottum | 0.071224 |
| stream_stream | fils | schiltach | 0.085424 |
| stream_stream | fils | wilde_gutach | 0.065468 |
| stream_stream | fils | wutach | 0.080822 |
| hatchery_hatchery | fischzucht_hofer | fischzucht_lohmuehle | 0.176091 |
| hatchery_hatchery | fischzucht_hofer | fischzucht_stoerk | 0.240611 |
| stream_hatchery | fischzucht_hofer | grosse | 0.216084 |
| stream_hatchery | fischzucht_hofer | gutach | 0.198185 |
| stream_hatchery | fischzucht_hofer | hoellenbach | 0.223 |
| stream_hatchery | fischzucht_hofer | insenbach | 0.280419 |
| stream_hatchery | fischzucht_hofer | kinzig | 0.162565 |
| stream_hatchery | fischzucht_hofer | kleine_Wiese | 0.191201 |
| stream_hatchery | fischzucht_hofer | lauter | 0.241455 |
| stream_hatchery | fischzucht_hofer | lautracher_Ach | 0.216959 |
| stream_hatchery | fischzucht_hofer | rottum | 0.165649 |
| stream_hatchery | fischzucht_hofer | schiltach | 0.186759 |
| stream_hatchery | fischzucht_hofer | wilde_gutach | 0.153805 |
| stream_hatchery | fischzucht_hofer | wutach | 0.166422 |
| hatchery_hatchery | fischzucht_lonmuehle | fischzucht_stoerk | 0.172857 |
| stream_hatchery | fischzucht_lohmuehle | grosse | 0.152067 |
| stream_hatchery | fischzucht_lohmuehle | gutach | 0.129972 |
| stream_hatchery | fischzucht_lohmuehle | hoellenbach | 0.168231 |
| stream_hatchery | fischzucht_lohmuehle | insenbach | 0.227137 |
| stream_hatchery | fischzucht_lohmuehle | kinzig | 0.118995 |
| stream_hatchery | fischzucht_lohmuehle | kleine_Wiese | 0.124342 |
| stream_hatchery | fischzucht_lohmuehle | lauter | 0.191601 |
| stream_hatchery | fischzucht_lohmuehle | lautracher_Ach | 0.172534 |
| stream_hatchery | fischzucht_lohmuehle | rottum | 0.10573 |
| stream_hatchery | fischzucht_lohmuehle | schiltach | 0.127996 |
| stream_hatchery | fischzucht_lohmuehle | wilde_gutach | 0.101682 |
| stream_hatchery | fischzucht_lohmuehle | wutach | 0.119645 |
| stream_hatchery | fischzucht_stoerk | grosse | 0.183555 |
| stream_hatchery | fischzucht_stoerk | gutach | 0.165088 |
| stream_hatchery | fischzucht_stoerk | hoellenbach | 0.201559 |
| stream_hatchery | fischzucht_stoerk | insenbach | 0.254308 |
| stream_hatchery | fischzucht_stoerk | kinzig | 0.15464 |
| stream_hatchery | fischzucht_stoerk | kleine_Wiese | 0.1525 |
| stream_hatchery | fischzucht_stoerk | lauter | 0.230755 |
| stream_hatchery | fischzucht_stoerk | lautracher_Ach | 0.210144 |
| stream_hatchery | fischzucht_stoerk | rottum | 0.137749 |
| stream_hatchery | fischzucht_stoerk | schiltach | 0.15566 |

(Table S1 continued)

| Comparison | Pop1 | Pop2 | Weighted_Fst |
|---|---|---|---|
| stream_hatchery | fischzucht_stoerk | wilde_gutach | 0.144109 |
| stream_hatchery | fischzucht_stoerk | wutach | 0.157219 |
| stream_stream | grosse | gutach | 0.142761 |
| stream_stream | grosse | hoellenbach | 0.17907 |
| stream_stream | grosse | insenbach | 0.228423 |
| stream_stream | grosse | kinzig | 0.131789 |
| stream_stream | grosse | kleine_Wiese | 0.133576 |
| stream_stream | grosse | lautracher_Ach | 0.179119 |
| stream_stream | grosse | rottum | 0.117982 |
| stream_stream | grosse | schiltach | 0.140915 |
| stream_stream | grosse | wilde_gutach | 0.114529 |
| stream_stream | grosse | wutach | 0.129668 |
| stream_stream | gutach | hoellenbach | 0.158621 |
| stream_stream | gutach | insenbach | 0.210854 |
| stream_stream | gutach | kinzig | 0.115367 |
| stream_stream | gutach | kleine_Wiese | 0.115165 |
| stream_stream | gutach | lauter | 0.178625 |
| stream_stream | gutach | lautracher_Ach | 0.161943 |
| stream_stream | gutach | rottum | 0.103385 |
| stream_stream | gutach | schiltach | 0.119359 |
| stream_stream | gutach | wilde_gutach | 0.097439 |
| stream_stream | gutach | wutach | 0.113787 |
| stream_stream | hoellenbach | insenbach | 0.24413 |
| stream_stream | hoellenbach | kinzig | 0.14031 |
| stream_stream | hoellenbach | kleine_Wiese | 0.149687 |
| stream_stream | hoellenbach | lauter | 0.213873 |
| stream_stream | hoellenbach | lautracher_Ach | 0.195509 |
| stream_stream | hoellenbach | rottum | 0.136008 |
| stream_stream | hoellenbach | schiltach | 0.158277 |
| stream_stream | hoellenbach | wilde_gutach | 0.125423 |
| stream_stream | hoellenbach | wutach | 0.142911 |
| stream_stream | insenbach | kinzig | 0.193993 |
| stream_stream | insenbach | kleine_Wiese | 0.209836 |
| stream_stream | insenbach | lauter | 0.27724 |
| stream_stream | insenbach | lautracher_Ach | 0.250372 |
| stream_stream | insenbach | rottum | 0.191376 |
| stream_stream | insenbach | schiltach | 0.215875 |
| stream_stream | insenbach | wilde_gutach | 0.183016 |
| stream_stream | insenbach | wutach | 0.202309 |
| stream_stream | kinzig | kleine_Wiese | 0.108966 |
| stream_stream | kinzig | lauter | 0.160299 |
| stream_stream | kinzig | lautracher_Ach | 0.14587 |
| stream_stream | kinzig | rottum | 0.090287 |
| stream_stream | kinzig | schiltach | 0.100602 |
| stream_stream | kinzig | wilde_gutach | 0.079174 |
| stream_stream | kinzig | wutach | 0.096519 |
| stream_stream | kleine_Wiese | lauter | 0.171143 |
| stream_stream | kleine_Wiese | lautracher_Ach | 0.156145 |
| stream_stream | kleine_Wiese | rottum | 0.090304 |
| stream_stream | kleine_Wiese | schiltach | 0.109952 |
| stream_stream | kleine_Wiese | wilde_gutach | 0.089341 |
| stream_stream | kleine_Wiese | wutach | 0.104503 |

(Table S1 continued)

| Comparison | Pop1 | Pop2 | Weighted_Fst |
|---|---|---|---|
| stream_stream | lauter | lautracher_Ach | 0.005308 |
| stream_stream | lauter | rottum | 0.15617 |
| stream_stream | lauter | schiltach | 0.178913 |
| stream_stream | lauter | wilde_gutach | 0.139878 |
| stream_stream | lauter | wutach | 0.157529 |
| stream_stream | lautracher | Ach_rottum | 0.141843 |
| stream_stream | lautracher | Ach_schiltach | 0.161184 |
| stream_stream | lautracher | Ach_wilde_gutach | 0.126134 |
| stream_stream | lautracher | Ach_wutach | 0.142955 |
| stream_stream | rottum | schiltach | 0.092918 |
| stream_stream | rottum | wilde_gutach | 0.07431 |
| stream_stream | rottum | wutach | 0.09023 |
| stream_stream | schiltach | wilde_gutach | 0.092235 |
| stream_stream | schiltach | wutach | 0.104882 |
| stream_stream | wilde_gutach | wutach | 0.07612 |

**Table S2.** Output from genome-wide association study of 3 hatchery and 14 stream populations of rainbow trout uncovering positions in the genome that are associated with the origin (hatchery or stream). Headers: rs = SNP ID, chromo = chromosome, pos = position on chromosome in bp, n-miss = number of missing individuals for each site, allele 1 = genotype for allele 1, allele 2 = genotype for allele 2, af = allele frequency of minor allele, beta = bata estimates, se = standard error for beta, logl_H1 = log-likelihood under the alternative hypothesis, l_remle = remle estimates for lambda, p-wald = p-value from Wald test.

| rs | chromo | pos | ps | n_miss | allele1 | allele0 | af | beta | se | logl_H1 | l_remle | p_wald | Inside_gene | upstream_gene | downstream_gene |
|---|---|---|---|---|---|---|---|---|---|---|---|---|---|---|---|
| NC_048569.1_71720723 | NC_048569 | 71720723 | -9 | 0 | T | C | 0.075 | -0.2235376 | 0.03543668 | 102.4476 | 100000 | 1.56E-09 | NA | faslg | fam20b |
| NC_048569.1_73113400 | NC_048569 | 73113400 | -9 | 2 | A | C | 0.067 | -0.2716197 | 0.04048436 | 104.6552 | 100000 | 1.68E-10 | LOC110524521 | mdp1 | LOC110524520 |
| NC_048569.1_73113477 | NC_048569 | 73113477 | -9 | 5 | T | G | 0.067 | -0.2470505 | 0.03849006 | 103.0595 | 100000 | 8.53E-10 | LOC110524521 | mdp1 | LOC110524520 |
| NC_048569.1_73188394 | NC_048569 | 73188394 | -9 | 0 | T | C | 0.057 | -0.2816353 | 0.04037755 | 106.1621 | 100000 | 3.64E-11 | LOC110522885 | LOC110524524 | LOC110524525 |
| NC_048574.1_65703509 | NC_048574 | 65703509 | -9 | 2 | A | C | 0.059 | -0.2347409 | 0.03786246 | 101.8017 | 100000 | 2.81E-09 | LOC110534492 | LOC110534493 | LOC110534495 |
| NC_048574.1_65743274 | NC_048574 | 65743274 | -9 | 0 | T | C | 0.062 | -0.2503512 | 0.03509779 | 106.9723 | 100000 | 1.45E-11 | NA | LOC110534492 | LOC110534495 |
| NC_048575.1_45024635 | NC_048575 | 45024635 | -9 | 0 | A | C | 0.938 | 0.2521843 | 0.04195454 | 101.0285 | 100000 | 7.72E-09 | NA | LOC110535028 | LOC110535732 |
| NC_048577.1_37409778 | NC_048577 | 37409778 | -9 | 7 | T | A | 0.643 | 0.1210512 | 0.02008176 | 101.0941 | 100000 | 7.06E-09 | rbtstat3 | stat5 | LOC110486263 |
